# A new product containing secretome purified from bovine colostrum, Purasomes IRC100+, in the treatment of vaginal atrophy: *in vitro* study to evaluate safety and effectiveness

**DOI:** 10.1101/2024.12.31.630871

**Authors:** Antonio Salvaggio, Anna Privitera, Stefania Pennisi, Maria Violetta Brundo

**Affiliations:** Department of Biological, Geological and Environmental Science, University of Catania, Catania, Italy

**Keywords:** vaginal atrophy, MTT test, scratch-wound assay, VK2/E6E7 cell line, fibroblasts

## Abstract

Vaginal atrophy consists of the loss of elasticity and thinning of the vaginal tissues and mucous membranes. This is a condition that affects women especially during menopause, following the hormonal decline resulting from the cessation of ovarian activity. The objective of the present study was evaluated the effects of Purasomes IRC100+ on the proliferation and migration of VK2/E6E7 vaginal epithelial cells and on fibroblasts by *in vitro* experiments. We tested three different concentrations (0.5%, 1% and 2%). We found that cell viability was highest at 2% Purasomes IRC100+ concentration for both cell lines. The scratch-wound assay was also done. Compared with the cell scratch at 0 h, cells in all groups migrated after 24 h, the scratch was significantly reduced in treated cells compared to untreated cells. In particular, the cell migration rates were 77% and 82%, respectively for VK2/E6E7 cells and fibroblasts treated with 2% of Purasomes IRC100+. Finally, this product promotes vaginal epithelial cell proliferation and therefore it could improve vaginal atrophy and vaginal wound healing.

## 1. Introduction

Vaginal atrophy is a very common pathology in women, especially during menopause. Vaginal atrophy has a prevalence of 65% after one year after menopause reaching values of 90% after 20 years after menopause. The incidence of this pathology is certainly underestimated, as emerges from numerous studies [1-6]. A study conducted in Europe on 4201 women and aimed to investigate women’s opinions, attitudes and perceptions on menopause in general and on treatments for menopause symptoms has highlighted that European women need and want to be better informed and educated on the implications of vaginal atrophy on their quality of life [1]. But the most worrying aspect emerged from another North American study which shows that only around 25% of women who suffer from them spontaneously communicate these problems to their doctor out of modesty or reluctance, and 70% of those interviewed report that only rarely (if ever!) does their doctor ask them questions about problems such as vaginal dryness [2]. This situation has not changed in recent years as emerges from various studies [3-6]. Estrogen receptors are present on the vaginal epithelium which determine the maintenance of the elasticity and thickness of the mucous layer and vaginal walls. During menopause, due to their reduction, the epithelium becomes less vascularized, thinner and glycogen also decreases. Furthermore, the amount of collagen in the connective tissue, which has the function of supporting the epithelium, also decreases, and this determines the loss of the normal roughness of the inside of the vagina. All these factors contribute to the loss of the protective function of the mucosa and therefore there is greater susceptibility [7,8]. In addition to menopause, a drop in estrogen can also occur in other periods, for example during breastfeeding, after removal of the ovaries (surgical menopause), after chemotherapy, after pelvic radiotherapy, after hormone therapy for cancer at the breast [9,10].

Aims of this study was to evaluate the effects of a new products Purasomes IRC100+, that activating collagen and elastin at a molecular level [11,12], restores all vaginal functions such as secretion, absorption, elasticity, lubrication and vaginal epithelium thickness. This product containing bovine colostrum secretomes uses a new technology, called AMPLEX plus, that contains totipotent bioactive elements purified from bovine colostrum, such as growth factors and cytokines, and exosomes passively loaded with these actives’ biomolecules. The secretome including proteins and other biomolecules secreted or released by a cell or tissue into the extracellular environment [13]. They play a pivotal role in facilitating intercellular communication, regulating immune responses and tissue repair [13]. Exosomes are extracellular vesicles that have an important role in cellular communication and tissue repair [14]. Exosomes act synergistically with secretome to promote multiple signaling pathways [14]. Recent research evidenced the secretome from colostrum bovine to have a therapeutic potential high [15,16]. In particular, growth factors present in secretome and in exosomes, such as bFGF, have a promoting effect on vaginal wound healing *in vitro*, because promotes proliferation, cell differentiation and collagen production in fibroblast [12].

## 2. Materials and Methods

For our research we purchased a commercial product (Purasomes IRC100+) that utilize new technologies containing among the ingredient’s colostrum bovine secretome. In particular, the product contains exosomes purified from bovine colostrum passively loaded with growth factors and cytokines purified from bovine colostrum. This latest technology is called AMPLEX plus technology.

### 2.1 Propagation and maintenance of VK2-E6E7 cells

The VK2-E6E7, normal human vaginal cells (ATCC CRL2616), was maintained and propagated according to the ATCC protocol using keratinocyte serum-free medium (KSFM) supplemented with 50 mg/mL^-1^ bovine pituitary extract and 0.1 ng/mL^-1^ epidermal growth factor. The cells were grown at 37°C with 5% CO_2_ and 100% humidity and split every 3-4 days depending on cell confluence. To prevent bacterial contamination of the media, 100 IU mL^-1^ of penicillin and streptomycin was added.

### 2.2 Propagation and maintenance of fibroblasts

Human non immortalized fibroblasts were cultured in Dulbecco’s Modified Eagle Medium (DMEM) supplemented with 10% FBS, streptomycin (0.3 mg mL^-1^), penicillin (50 IU mL^-1^) and GlutaMAX (1 mM) by using 25 or 75 cm^2^ polystyrene culture flasks. Cells were maintained in a humidified environment (37 °C, 95% air/5% CO_2_), and split every 2–3 days depending on cell confluence.

### 2.3 Analysis of cell proliferation

Both cell lines were treated with Purasomes IRC100+ (0.5% and 1%) and incubated for 24 hours in a humidified environment (37°C, 5% CO_2_). At the end of the treatment, cell proliferation/metabolic activity were measured by the well-known MTT assay. Briefly, at the end of the treatment, MTT solution (1 mg/mL in DMEM medium) was added to each well and cells were incubated for 2 hours in a humidified environment (37°C, 5% CO_2_). During the final step, DMSO was used to melt the crystals, while the Synergy H1 Hybrid Multi-Mode Microplate Reader (Biotek, Shoreline, WA, USA) was used to read the absorbance at 569 nm. Values were normalized with respect to control untreated cells (VK2-E6E7 cells and fibroblasts) and were expressed as the percent variation of cell proliferation/metabolic activity.

### 2.4 Scratch-wound assay

VK2/E6E7 cells and fibroblasts were inoculated separately into a 6-well culture plate and the cell monolayer was scratched using a sterile 200 µl pipette tip. Cells were washed three times with sterile PBS to remove scratched cells, and serum-free medium was added. Purasomes IRC100+ were added in addition to the blank control group and cultured continuously for 24 hours. We tested three different concentrations (0.5%, 1% and 2%) and all experiments were replicated three times.

The wound closure was calculated using the equation:

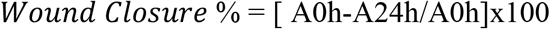

where A0h is the area (pixels) of the wound calculated after scratching (t = 0 h) and A24h is the area (pixels) of the unhealed wound (which is not covered by the cells) that remained 24 h after the scratching.

### 2.5 Statistical analysis

The statistical analysis was carried out by using Graphpad Prism software, version 8.0 (Graphpad software, San Diego, CA, USA). Two-way analysis of variance (ANOVA), followed by Tukey’s post hoc test, was used for multiple comparisons. The statistical significance was set at p-values < 0.01. Data were reported as the mean ± SD of at least three independent experiments.

## 3. Results and Discussions

Reproductive aging has several implications on women’s health, including vaginal atrophy that represents a primary manifestation of reproductive aging. A decrease in estrogen levels, in fact, it results in a fragile and thin vaginal epithelium. This condition also leads to diminished local blood flow and an increase in vaginal pH, adversely affecting quality of life for menopausal women [17, 18]. Although estrogen therapy can currently improve symptoms of vaginal atrophy, its long-term use is controversial due to potential risks for women ‘s health [19, 20]. On the vaginal epithelium there are estrogen receptors which, by binding to them, activate them and determine the maintenance of the elasticity and thickness of the mucous layer and vaginal walls. With their reduction or even disappearance, the epithelium becomes less vascularized, thinner and glycogen also decreases. Furthermore, the amount of collagen in the connective tissue, which has the function of supporting the epithelium, also decreases, and this determines the loss of the normal roughness of the inside of the vagina. All these factors are fundamental for the protection of the vaginal mucosa which becomes very susceptible to possible trauma [17, 18]. The treatment of vaginal atrophy should in fact have as its objectives the improvement and relief of vaginal symptoms and the recovery of vaginal physiology with improvement of epithelial trophism. The treatments available today are of two types: non-hormonal (such as phytoestrogens, moisturizers and vitamins) and hormonal. Phyto-estrogens are non-steroidal molecules of plant origin derived from soy and red clover, which bind to estrogen receptors exerting an estrogen-like effect. Several studies have shown that the use of red clover isoflavones reduces parabasal cells and increases superficial cells, thus increasing the vaginal maturation index, without exerting a significant effect on endometrial thickness [21] but had a positive effect on triglyceride levels [22], however always demonstrating poor efficacy in the treatment of dystrophic symptoms of menopause [21, 22]. Furthermore, the risk/benefit ratio of phyto-estrogens with breast cancer is still debated [23]. Moisturizing products are complex polymers that behave like bioadhesives that adhere to the epithelial cells of the vaginal wall and to the mucins, retaining water. The positive effect on the symptoms of vaginal atrophy is related to the fact that these products have the property of tampons, resulting in a reduction in vaginal pH [24]. Systemic hormone therapy involves the administration of estrogens associated with progestins in non-hysterectomized patients, while estrogen alone in those who have undergone hysterectomy. It is administered according to two therapeutic schemes: one continuous and one sequential. The contraindications to the use of hormone replacement therapy are various, including previous breast or endometrial cancer, alterations in liver function, previous venous thrombosis or functional events and the presence of atypical blood losses [25]. It is also possible to carry out local therapy with vaginal estrogen, which is certainly more effective in treating vaginal problems and at the moment does not seem to have the systemic negative effects of systemic hormone therapy [26]. These benefits are lost once hormone therapy is stopped [26]. Today it is also possible to carry out laser treatments for vaginal rejuvenation. The use of laser allows you to restore the structure of the vaginal mucosa up to the pre-menopausal period [27], but to date its use is not always recommended [28, 29]. Several Authors believe that an adequately selected treatment could lead to restoration and maintenance of the vaginal function and vaginal health.

We evaluated the effects of Purasomes IRC100+ on the proliferation and migration of VK2/E6E7 vaginal epithelial cells and on fibroblasts by *in vitro* experiments. We found that cell viability was highest at 2% Purasomes IRC100+ concentration for both VK2/E6E7 cells (Figure 1) and fibroblasts (Figure 2). The migration of VK2/E6E7 cells and fibroblasts by Purasomes IRC100+ were also explored. Compared with the cell scratch at 0 h, cells in all groups migrated after 24 h, the scratch was significantly reduced in treated cells compared to untreated cells (Figs 3, 4). In particular, the cell migration rates were 77% and 25%, respectively for VK2/E6E7 cells and untreated cells (Figure 3); the cell migration rates were 82% and 24%, respectively for fibroblasts and untreated cells (Figure 4).

**Figure 1.**
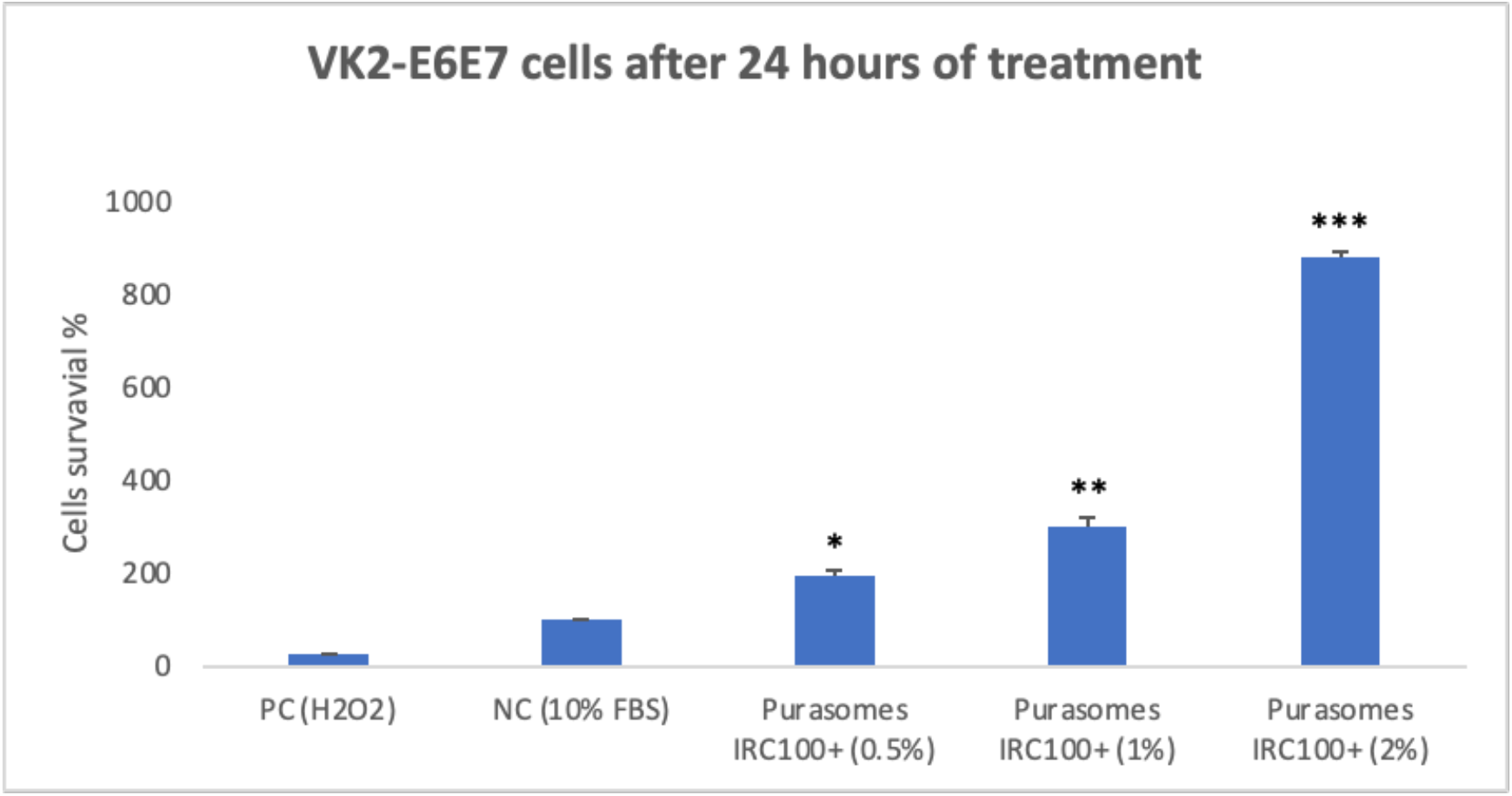
The cell survival of normal human vaginal cells (VK2-E6E7) at 24h following treatment with Purasomes IRC100+ to different concentrations (0.5%, 1% and 2%). NC (Negative Control, cells without treatment); PC (Positive Control, 80 µM H_2_O_2_); The asterisks denote the degree of significance between results: *p<0.01; **p<0.001; ***p<0.0001. Errors bars represent the Standard Deviation of the mean (experiment was repeated three times).

**Figure 2.**
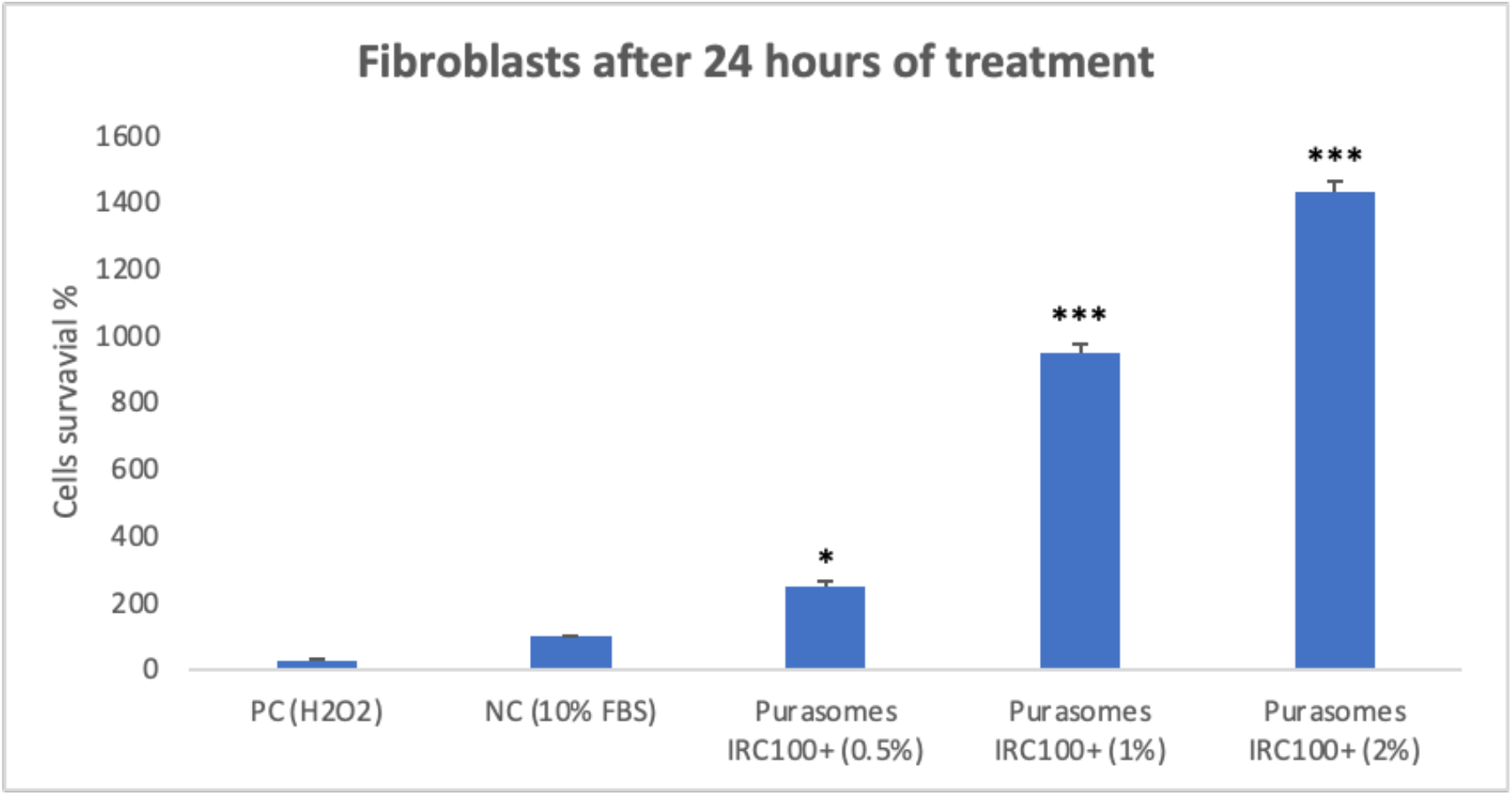
The cell survival of fibroblasts at 24h following treatment with Purasomes IRC100+ to different concentrations (0.5%, 1% and 2%). NC (Negative Control, cells without treatment); PC (Positive Control, 80 µM H_2_O_2_); The asterisks denote the degree of significance between results: *p<0.01; ***p<0.0001. Errors bars represent the Standard Deviation of the mean (experiment was repeated three times).

**Figure 3.**
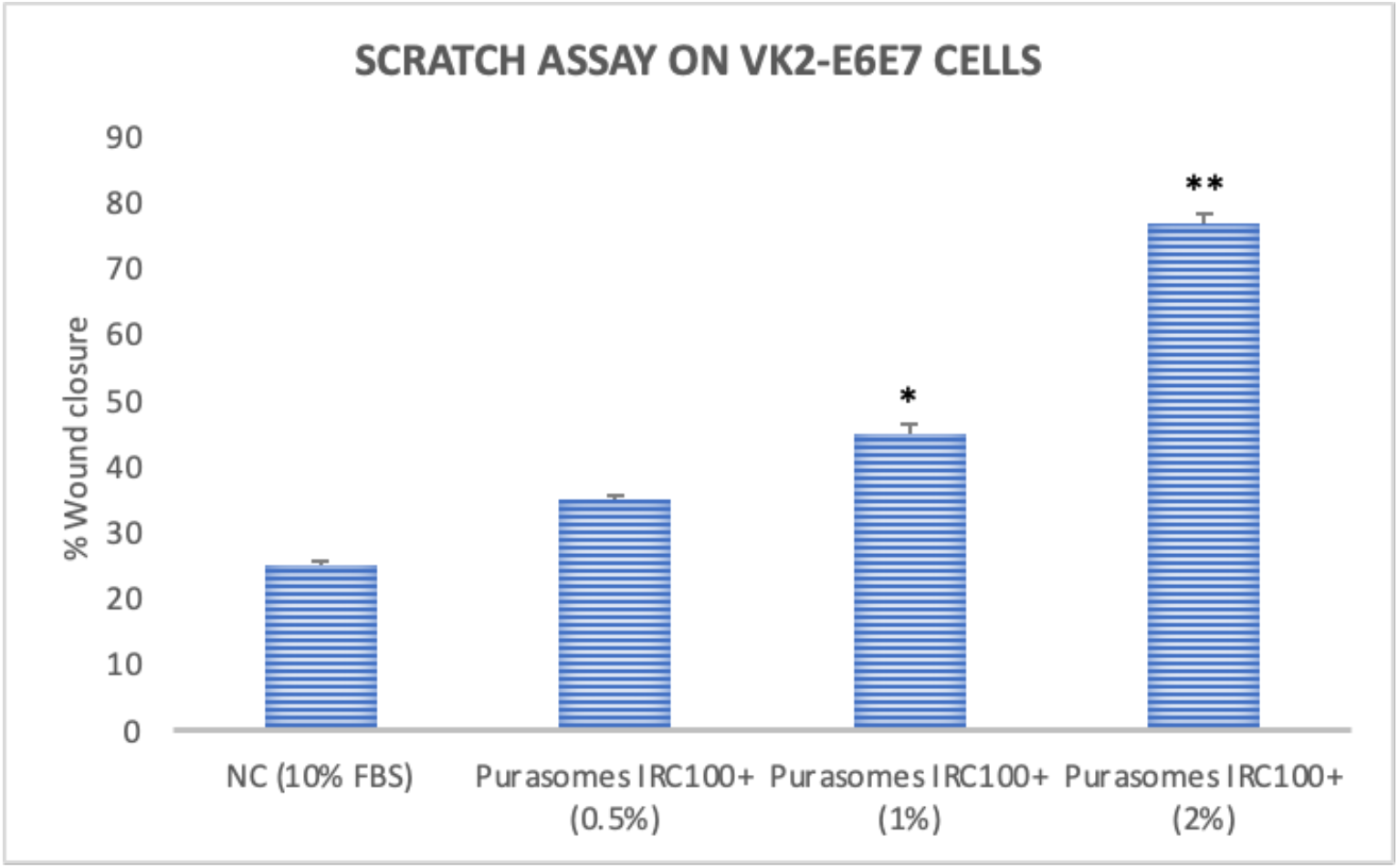
Scratch assay on of normal human vaginal cells (VK2-E6E7) at 24 hours’ time. Untreated cells were considered as a control (NC). The asterisks denote the degree of significance between results: *p<0.01; **p<0.001. Errors bars represent the Standard Deviation of the mean (experiment was repeated 3 times).

**Figure 4.**
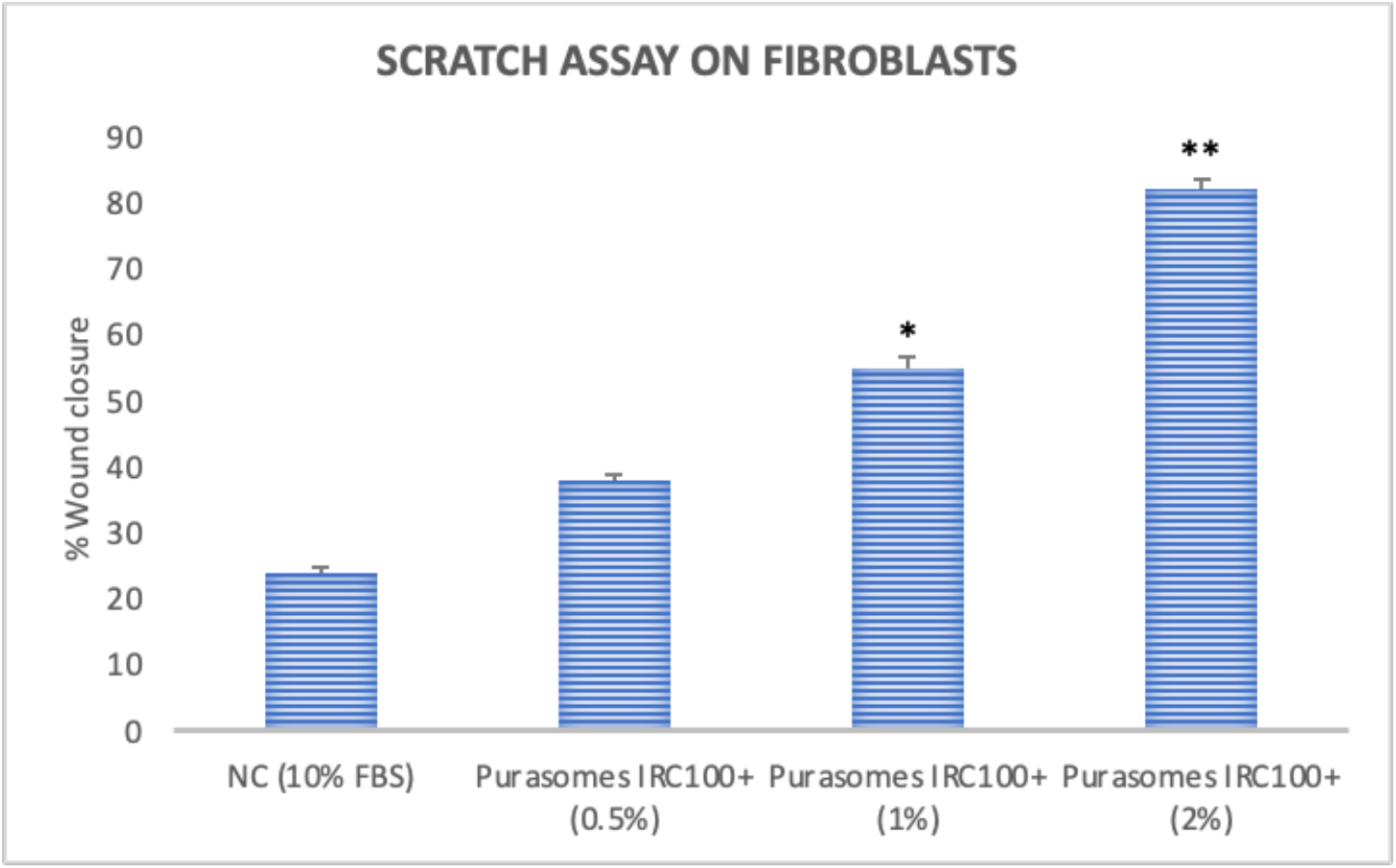
Scratch assay on of human fibroblasts at 24 hours’ time. Untreated cells were considered as a control (NC). The asterisks denote the degree of significance between results: *p<0.01; **p<0.001. Errors bars represent the Standard Deviation of the mean (experiment was repeated 3 times).

The product tested, contains the bovine colostrum secretome, that due to their rich composition maximize healing processes and accelerate tissue repair [30, 31]. In particular, it contains purified and concentrated exosomes, passively loaded with numerous biomolecules purified from bovine colostrum. Colostrum is the first secretion of mammalian glands, and its components, biologically active, have beneficial effects. Bovine colostrum has the highest concentration of biomolecules (such as growth factors and cytokines) and could provide health benefits. The growth factors present in bovine colostrum can enhance elasticity and hydration of vagina mucosa [32]. Several studies show that growth factors, particularly basic fibroblast growth factor and epidermal growth factor, can promote vaginal wound healing *in vitro*. It has been proven that basic fibroblast growth factor promotes proliferation, differentiation, and collagen types I and III production. Epidermal growth factor stimulates proliferation and connective tissue growth factor promotes Tenascin-C expression [33-35]. Exosomes are nanoscale-sized vesicles that are released by almost all eukaryotic cells, including the bovine mammary gland [36-38]. The double lipid layer of exosomes is enriched in lipid components such as cholesterol and sphingolipids, which are responsible for the stability of exosomes and their ability to interact with target cell membranes. Within the lipid bilayer, there are also specific membrane proteins, including tetraspanins (CD9, CD63, and CD81), which have a role in the fusion of exosomes with the membranes of other cells. They are essential in intercellular communication, and their cargo is extremely heterogenous [38]. Exosomes from bovine colostrum have been shown to have great tissue repair potential, particularly in moderating the effects of skin ageing [39-43]. This effect is mediated, in part, by activating the Wnt/β-catenin signaling pathway, which lead to increased production of collagen, responsible for skin elasticity and strength, but also from the reduction the level of metalloproteinases enzymes that have an important role in tissue remodeling and, their excessive activity, can lead to premature skin ageing [42-43]. In addition, exosomes significantly can reduce intracellular oxidative stress in keratinocytes exposed to UV-C radiation [40]. Exosomes derived from bovine colostrum also demonstrate the potential to improve skin hydration and reduce wrinkles [44].

## 4. Conclusions

In conclusion, we showed that Purasomes IRC100+ promotes vaginal epithelial cell proliferation, vaginal wound healing and appears to be a promising formulation to improve the trophicity and hydration of the vaginal mucosa in the treatment of vaginal atrophy. Treatment with this new product could lead to restoration and maintenance of the vaginal function, solving the problems not only of menopausal women, but also all problems that young women have during breastfeeding, after removal of the ovaries, after chemotherapy, after pelvic radiotherapy, after hormone therapy for breast cancer.

## Author Contributions

Conceptualization, A.S. and M.V.B.; methodology, A.P. and S.P.; software, A.P.; validation, A.P.; formal analysis, S.P.; data curation, A.S..; writing-original draft preparation, A.S. and M.V.B.; writing-review and editing, A.S., and M.V.B.; supervision, A.S. and M.V.B.; funding acquisition, M.V.B. All authors have read and agreed to the published version of the manuscript.

## Institutional Review Board Statement

This study was performed in line with the principles of the Declaration of Helsinki and does not require approval by the Ethics Committee of University of Catania.

## Informed Consent Statement

Not applicable.

## Data Availability Statement

The data presented in this study are available on request from the corresponding author.

## Acknowledgments

S.P. thanks the Ph.D. program.

